# PU.1 expression defines distinct functional activities in the phenotypic HSC compartment in a mouse model of inflammatory stress

**DOI:** 10.1101/2021.10.25.465758

**Authors:** James S. Chavez, Jennifer L. Rabe, Giovanny Hernandez, Taylor S. Mills, Katia E. Niño, Pavel Davizon-Castillo, Eric M. Pietras

**Affiliations:** Division of Hematology, University of Colorado Anschutz Medical Campus, Aurora, CO, 80045 USA; Department of Immunology and Microbiology, University of Colorado Anschutz Medical Campus, Aurora, CO, 80045 USA; Department of Pediatrics, University of Colorado Anschutz Medical Campus, Aurora, CO, 80045 USA

## Abstract

The transcription factor PU.1 is a critical regulator of lineage fate in blood-forming hematopoietic stem cells (HSC). In response to inflammatory signals, PU.1 expression is increased in HSC, activating myeloid differentiation genes while repressing cell cycle and protein synthesis genes. To address potential functional heterogeneity arising in the phenotypic HSC compartment due to changes in PU.1 expression, here we fractionated phenotypic HSC using the SLAM code in conjunction with PU.1 expression levels using the *PU*.*1-EYFP* reporter mouse strain. While PU.1^lo^ SLAM cells contain extensive long-term repopulating activity and a molecular signature corresponding to HSC activity at steady state, under inflammatory conditions the PU.1^lo^ SLAM fraction is comprised almost entirely of HSC-like cells containing extensive short-term megakaryocytic potential. Our data demonstrate that the phenotypic HSC gate is heterogenous, and that similar PU.1 transcription factor levels can be tied to distinct functional activities under steady-state and inflammatory conditions.

## Introduction

PU.1/*Spi1* is an Ets-family transcription factor that serves as a ‘master regulator’ of hematopoietic myeloid and lymphoid lineage fates^1^. Levels of PU.1 expression are lineage-specific, with the highest levels expressed in myeloid-committed cells whereas intermediate PU.1 levels are found in lymphocytes^2, 3^. On the other hand, megakaryocyte/erythroid committed cells express little to no PU.1^4^. In a context-dependent fashion, PU.1 can transactivate or repress a wide range of genes associated with lineage specification as well as numerous genes regulating cell cycle and metabolic activity^5-7^. The immature hematopoietic stem cell (HSC) compartment is generally thought to express very low levels of PU.1, in line with classical models of hematopoiesis describing HSC as truly immature cells that are not lineage-committed^8^. Nonetheless, PU.1 is required for normal HSC self-renewal activity and cell cycle regulation, and PU.1 deficiency leads to defects in myeloid and lymphoid lineage specification^9^. In response to inflammatory cytokines such as interleukin-1 (IL-1), PU.1 expression rapidly increases within the HSC compartment, leading to concurrent activation of myeloid lineage genes and repression of cell cycle and protein synthesis genes^6, 10-14^. Hence, inflammatory signals can ‘poise’ HSC to replenish myeloid lineage cells in response to injury or infection, while also restraining HSC proliferation to maintain a homeostatic pool size in the bone marrow (BM).

In murine models, transplantable HSC activity is prospectively enriched within the phenotypic Lineage^-^/c-Kit^+^/Flk2^-^/Sca-1^+^/CD48^-^/CD150^+^ (SLAM) fraction^15^. Using an extensively validated *PU*.*1-EYFP* fusion protein reporter mouse strain^6, 11, 13, 16^, we previously sub-fractionated the phenotypic SLAM compartment based on PU.1-EYFP levels to assess differences in cell cycle and protein synthesis gene expression at steady state and in response to acute and chronic IL-1 stimulation^6^. Relative to PU.1^lo^ SLAM cells, PU.1^hi^ SLAM cells expressed reduced levels of cell cycle genes including *Myc* and *Ccnd1* at steady state. Notably, these differences were exacerbated by IL-1 exposure. Strikingly, expression of HSC-enriched genes such as *Fgd5* and *Rarb* was significantly reduced in PU.1^lo^ SLAM cells following IL-1 exposure, coinciding with increased cell cycle activity specifically in the PU.1^lo^ fraction^6^.

These data prompted us to hypothesize that while long-term repopulating HSC (HSC^LT^) are classically considered to express very low levels of PU.1 owing to their immature state, the PU.1^lo^ SLAM fraction may contain functionally heterogenous activities. Here, we assess 1) the functional and molecular properties of PU.1^hi^ and PU.1^lo^ SLAM cells at steady state and in response to IL-1-mediated inflammation; and 2) the extent to which low PU.1 levels represent distinct hematopoietic activities within the phenotypic SLAM gate.

## Results

### PU.1 levels define distinct repopulating activities in the SLAM compartment

To better understand the association between PU.1 expression and HSC function within the SLAM compartment following chronic IL-1 exposure, we injected *PU*.*1-EYFP* reporter mice with IL-1 or PBS daily for 20 days to simulate a chronic inflammatory state (**Fig. 1A**)^11^. This treatment led to significant increases in peripheral blood myeloid and platelet counts, suggesting IL-1 stimulates their production by HSC^11^ (**Fig. S1A**). Thus, we next measured PU.1-EYFP expression in phenotypic long-term HSC (HSC^LT^;EPCR^+^/CD34^-^ SLAM cells) and the remaining quadrants of the SLAM gate, fractionated based on EPCR and CD34 expression^10^. While mean PU.1 levels appeared similar among different SLAM fractions at steady state, IL-1 induced PU.1-EYFP expression in the HSC^LT^-enriched EPCR^+^ SLAM fractions (**Fig. 1B-C**). These data are consistent with our published findings showing that HSC express basally low levels of PU.1 relative to lineage committed hematopoietic progenitors^6, 10, 11^. On the other hand, PU.1 expression was not induced in EPCR^-^ cells, which coincidentally possess limited long-term repopulating activity^10^. These data suggested the response to IL-1 is likely heterogenous within the SLAM compartment, with PU.1 levels potentially separating distinct functional activities within this gate. Thus, we next fractionated SLAM cells based on PU.1-EYFP reporter expression. PU.1-EYFP levels within the SLAM gate followed a roughly normal distribution and as previously characterized, IL-1 exposure increased the proportion of PU.1^hi^ SLAM cells (**Fig. 1D-E**)^6, 13^. To identify functional heterogeneity tied to PU.1 expression, we subdivided the SLAM compartment based on the upper and lower quartiles of PU.1-EYFP expression (**Fig. 1D-E**) and analyzed the functional properties of these fractions via myeloid colony forming unit (CFU) assay and transplantation into lethally irradiated recipients (**Fig. 1D**). Surprisingly, while the PU.1^lo^ SLAM fraction of PBS-treated mice exhibited robust clonogenic potential, colony forming activity was nearly absent in this fraction following IL-1 exposure (**Fig. 1F**). Likewise, we detected little to no myeloid/lymphoid donor chimerism from IL-1-exposed PU.1^lo^ SLAM cells in the peripheral blood, or donor-derived HSC in the BM (**Fig. 1G-I**). What little donor-derived chimerism could be detected from IL-1-exposed PU.1^lo^ SLAM cells exhibited a myeloid bias relative to donor output from IL-1-exposed PU.1^hi^ SLAM cells (**Fig. S1B-C**). On the other hand, PU.1^hi^ SLAM fractions exhibited extensive myeloid colony forming capacity and an intermediate long-term repopulating activity relative to PU.1^lo^ controls, with no evidence of myeloid bias (**Fig. 1H-I, Fig. S1B-C**). Taken together, these data suggest distinct functional activities are contained within the PU.1^lo^ SLAM gate and are separated by chronic IL-1 stimulation.

**Figure 1.**
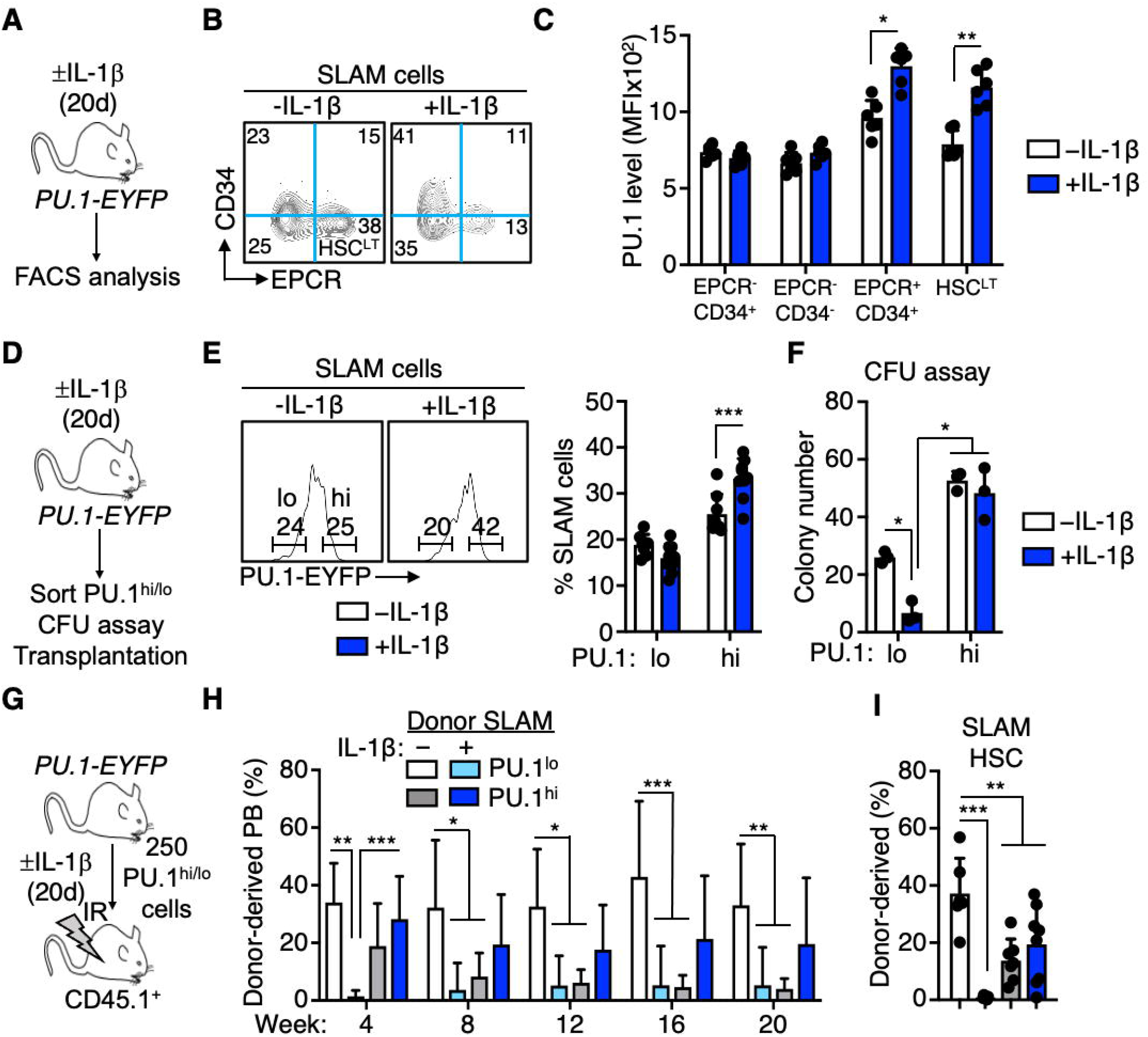
PU.1 levels define distinct repopulating activities in the SLAM compartment. **A)** Experimental design for analysis of *PU*.*1-EYFP* mice treated for 20d ± IL-1 (n = 5-6/grp). **B)** Representative FACS plot showing fractionation of SLAM gate by EPCR and CD34 levels. **C)** Analysis of *PU*.*1-EYFP* reporter expression in SLAM fractions. Data are expressed as geometric mean fluorescence intensity (MFI). Individual values are shown, bars represent mean values. Data are representative of two independent experiments. **D)** Experimental design for isolation and functional characterization of PU.1^hi/lo^ SLAM fractions of *PU*.*1-EYFP* mice treated for 20d ± IL-1β. **E)** Representative FACS plot (left) and summary data (right) showing frequency of PU.1^lo^ and PU.1^hi^ SLAM cells in *PU*.*1-EYFP* mice treated for 20d ± IL-1β (n = 6-8/grp). Individual values are shown, bars represent mean values. Data are compiled from two independent experiments. **F)** Colony numbers from CFU assay of PU.1^lo^ and PU.1^hi^ SLAM cells in *PU*.*1-EYFP* mice treated for 20d ± IL-1β (n = 3/grp). Individual values are shown, bars represent mean values. Data are representative of two independent experiments. **G)** Experimental design for isolation and transplant of purified PU.1^hi^ and PU.1^lo^ SLAM cells from *PU*.*1-EYFP* mice treated for 20d ± IL-1β into lethally irradiated recipient mice. **H)** Summary data showing donor peripheral blood chimerism in transplanted recipient mice (n = 6-8/grp). Data are compiled from two independent experiments. Bars represent mean values. **I)** Summary data showing donor BM SLAM cell chimerism in recipient mice at 20 weeks post-transplant (n = 6-8/grp). Individual values are shown, bars represent mean values. Data are compiled from two independent experiments. Data are shown as means ± SD. Significance was determined by ANOVA with Tukey’s post-test. *p<0.05; **p<0.01; ***p<0.001.

### The PU.1^lo^ SLAM fraction enriches for HSC-like Mk progenitors following IL-1 exposure

Given PU.1 serves as a myeloid differentiation factor, we had originally anticipated that the PU.1^lo^ SLAM compartment would retain extensive HSC potential regardless of IL-1 stimulation. To better understand the basis for our observations, we analyzed the phenotypic and molecular characteristics of the PU.1^lo^ and PU.1^hi^ SLAM fractions under inflammatory stress. Since we previously reported that EPCR and CD34 faithfully enrich for long-term HSC activity in the SLAM compartment under inflammatory stress conditions^10^, we assessed the relative changes in phenotypic EPCR^+^/CD34^-^ HSC^LT^ in the PU.1^hi^ and PU.1^lo^ SLAM compartments of *PU*.*1-EYFP* mice treated ± IL-1 for 20 days (**Fig. 2A**). As we expected, the PU.1^lo^ SLAM fraction from PBS-treated mice contained abundant phenotypic EPCR^+^/CD34^-^ HSC^LT^ relative to PU.1^hi^ cells, consistent with the superior long-term engraftment of PU.1^lo^ SLAM cells from PBS-treated mice (**Fig. 2B-D**). However, phenotypic HSC^LT^ were depleted from the PU.1^lo^ SLAM compartment following IL-1 exposure (**Fig. 2B-D**). This effect was likely due to IL-1-induced increases in PU.1 expression that shifts HSC^LT^ out of the phenotypic PU.1^lo^ gate^6^ (**Fig. 1C**). Notably, PU.1^lo^ and PU.1^hi^ SLAM fractions also contained a parallel population of EPCR^-^ cells (**Fig. 2B, Fig. S2A**). As we previously observed megakaryocyte (Mk) lineage gene signatures and elevated CD41 expression in EPCR^-^ SLAM fractions^10^, we reasoned that the parallel EPCR^-^ population in the PU.1^lo^ gate could contain previously characterized CD41^+^ HSC-like Mk progenitors^17^. Thus, we assessed levels of CD41 in the PU.1^lo^ and PU.1^hi^ SLAM compartments following chronic IL-1 exposure. Consistent with our hypothesis, the frequency and expression level of CD41 increased dramatically in PU.1^lo^ SLAM cells, but not PU.1^hi^ SLAM cells, following IL-1 exposure (**Fig. 2E-F**). Conversely, Mac-1 expression, a PU.1 target gene which we previously found to be increased in EPCR^+^ SLAM cells following IL-1 stimulation^10^, increased exclusively in PU.1^hi^ SLAM cells (**Fig. 2G**). The increased frequency of CD41^+^ cells was unlikely to be a result of megakaryocytic progenitors (MkP) contaminating the SLAM gate, as several key phenotypic features such as CD34, CD48, CD41, Sca-1, c-Kit and scatter properties were substantially different between PU.1^lo^ EPCR^-^ SLAM cells and phenotypic MkP^18^, regardless of IL-1 exposure (**Fig. S2A-B**). As a complementary readout, we analyzed gene expression patterns in PU.1^lo/hi^ SLAM cells using a Fluidigm qRT-PCR array. Consistent with our immunophenotypic analyses and published gene expression analyses of these fractions^6, 10^, levels of genes typically enriched in HSC, such as *Hoxb5, Egr1, Ifitm1* and *Foxo3* were significantly reduced in PU.1^lo^ SLAM cells (**Fig. 2I**). On the other hand, *Ifitm1* and *Foxo3*, as well as *Cpt1a*, were increased in IL-1-exposed PU.1^hi^ SLAM cells, suggesting alterations in metabolic and transcriptional activity that may be related to enforcement of quiescence and/or adaptation to an inflammatory environment. On the other hand, consistent with our previous published results^6^, the IL-1-exposed PU.1^lo^ SLAM fraction expressed higher levels of Mk-lineage genes such as *Gata1* and *Gfi1b*^19^ whereas the PU.1^hi^ fraction exhibited a pattern of increased myeloid lineage-associated genes including *Cd34, Cebpe, Csfr3* (GCSFR) and *Tnfrsf1a* (TNFR1) that was further potentiated by IL-1 exposure (**Fig. 2J**). Taken together, these data indicate that during inflammatory stress, the phenotypic PU.1^lo^ SLAM compartment becomes enriched for cells that contain extensive Mk lineage priming at the molecular level, whereas IL-1 exposure triggers elevated levels of myeloid gene expression in PU.1^hi^ SLAM cells. These changes likely underlie increased platelet and myeloid cell production observed in the peripheral blood following IL-1 stimulation.

**Figure 2.**
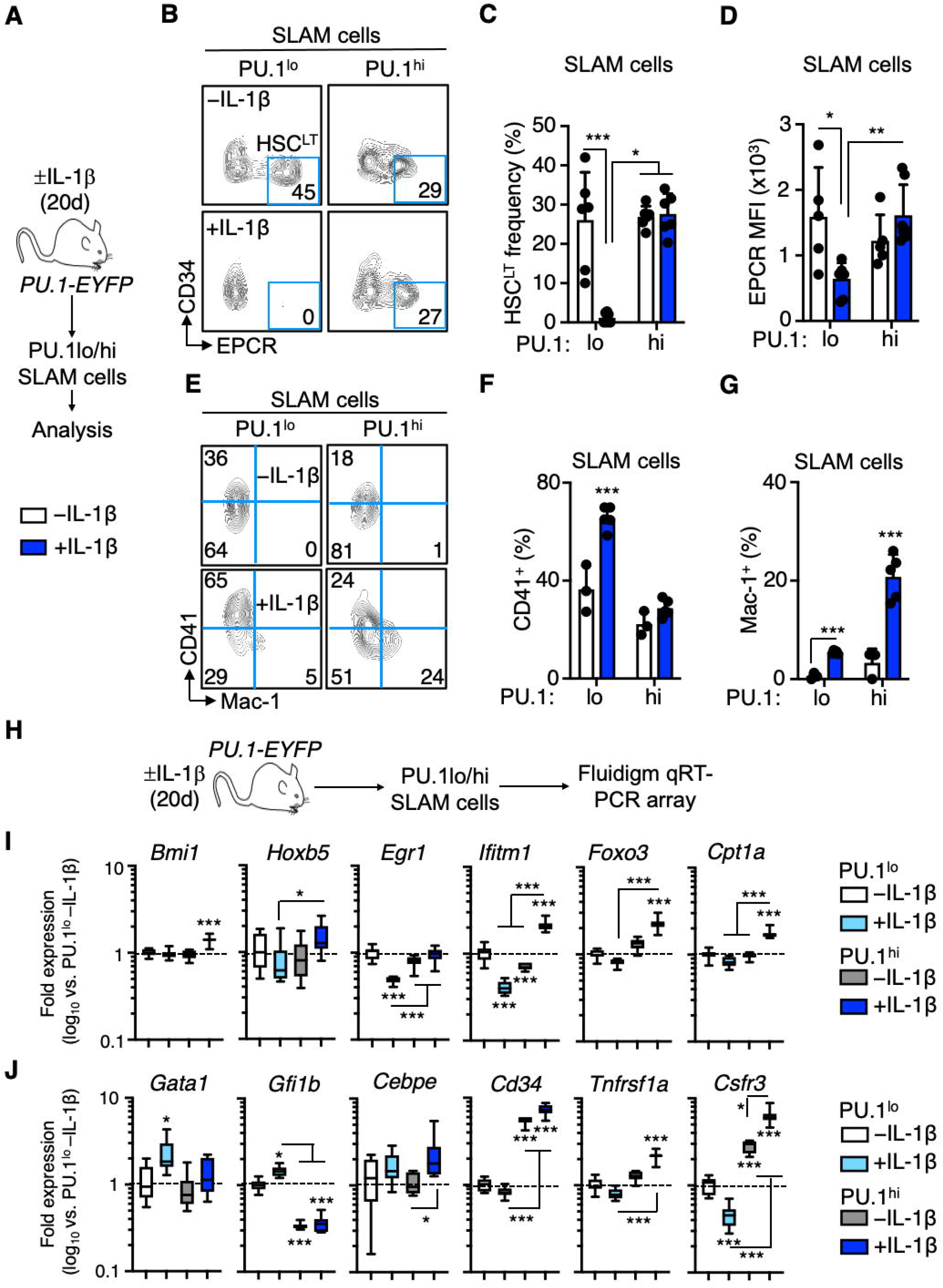
IL-1 exposure enriches for HSC-like Mk progenitors in the PU.1^lo^ SLAM fraction. **A)** Experimental design for analysis of SLAM cells from *PU*.*1-EYFP* mice treated for 20d ± IL-1β. **B)** Representative FACS plot showing phenotypic HSC^LT^ frequencies within the SLAM gate in *PU*.*1-EYFP* mice treated for 20d ± IL-1β. **C)** Summary data showing phenotypic HSC^LT^ frequency within the SLAM gate of *PU*.*1-EYFP* mice treated for 20d ± IL-1β (n = 6/grp). Individual values are shown, bars represent mean values. Data are compiled from two independent experiments. **D)** EPCR expression in phenotypic HSC^LT^ within the SLAM gate of *PU*.*1-EYFP* mice treated for 20d ± IL-1β (n = 6/grp). Data are expressed as geometric mean fluorescence intensity (MFI) Individual values are shown, bars represent mean values. Data are compiled from two independent experiments. **E)** Representative FACS plot showing frequencies of CD41^+^ and Mac-1^+^ cells within the SLAM gate in *PU*.*1-EYFP* mice treated for 20d ± IL-1β. **F)** Frequency of CD41+ cells within the SLAM gate of *PU*.*1-EYFP* mice treated for 20d ± IL-1β (n = 3-5/grp). Individual values are shown, bars represent mean values. Data are representative of two independent experiments. **G)** Frequency of Mac-1+ cells within the SLAM gate of *PU*.*1-EYFP* mice treated for 20d ± IL-1β (n = 3-5/grp). Individual values are shown, bars represent mean values. Data are representative of two independent experiments. **H)** Experimental design for Fluidigm qRT-PCR array analysis of PU.1^lo^ and PU.1^hi^ SLAM cells from *PU*.*1-EYFP* mice treated for 20d ± IL-1β (n = 8/grp). **I)** Fluidigm qRT-PCR analysis of self-renewal and metabolism genes in PU.1^lo^ and PU.1^hi^ SLAM cells from *PU*.*1-EYFP* mice treated for 20d ± IL-1β. Data are expressed as log_10_ fold change versus PU.1^lo^ -IL-1β. Box represents upper and lower quartiles with line representing median value. Whiskers represent minimum and maximum values. Data are representative of two independent experiments. **J)** Fluidigm qRT-PCR analysis of myeloid and megakaryocyte/erythroid genes in PU.1^lo^ and PU.1^hi^ SLAM cells from *PU*.*1-EYFP* mice treated for 20d ± IL-1β. Data are expressed as log_10_ fold change versus PU.1^lo^ -IL-1β. Box represents upper and lower quartiles with line representing median value. Whiskers represent minimum and maximum values. Data are representative of two independent experiments. Data are shown as means ± SD. Significance was determined by ANOVA with Tukey’s post-test. *p<0.05; **p<0.01; ***p<0.001.

### IL-1 rapidly triggers phenotypic remodeling of the PU.1^lo^ SLAM compartment

We next addressed whether the compositional changes in the PU.1^lo^ SLAM fraction are induced in response to acute inflammatory signals. We analyzed the SLAM compartment of PU.1-EYFP mice 24h (1 day) after a single injection of IL-1 or PBS (**Fig. 3A**). Similar to chronic IL-1 exposure, increased PU.1 expression was confined to EPCR^+^ SLAM cells including HSC^LT^. (**Fig. 3B-C**). Likewise, phenotypic HSC^LT^ were rapidly depleted in the PU.1^lo^ SLAM fraction, consistent with our published observation that PU-1-EYFP expression in these cells increases following IL-1 stimulation^6^ (**Fig. 3D-G)**. The frequency and expression of CD41 also rapidly increased in the PU.1^lo^ fraction (**Fig. 3H-J**). On the other hand, Mac-1 was not yet induced in PU.1^hi^ SLAM cells following acute IL-1 stimulation (Fig. 3J). Lastly, patterns of lineage-specific gene expression between PU.1^lo^ and PU.1^hi^ SLAM fractions were nearly identical to gene expression changes following chronic IL-1 treatment (**Fig. 3K-L**). Collectively, these data demonstrate that compositional changes within the SLAM compartment are rapidly established in response to inflammation, with the PU.1^lo^ SLAM fraction almost exclusively harboring EPCR^-^ cells that express Mk lineage markers.

**Figure 3:**
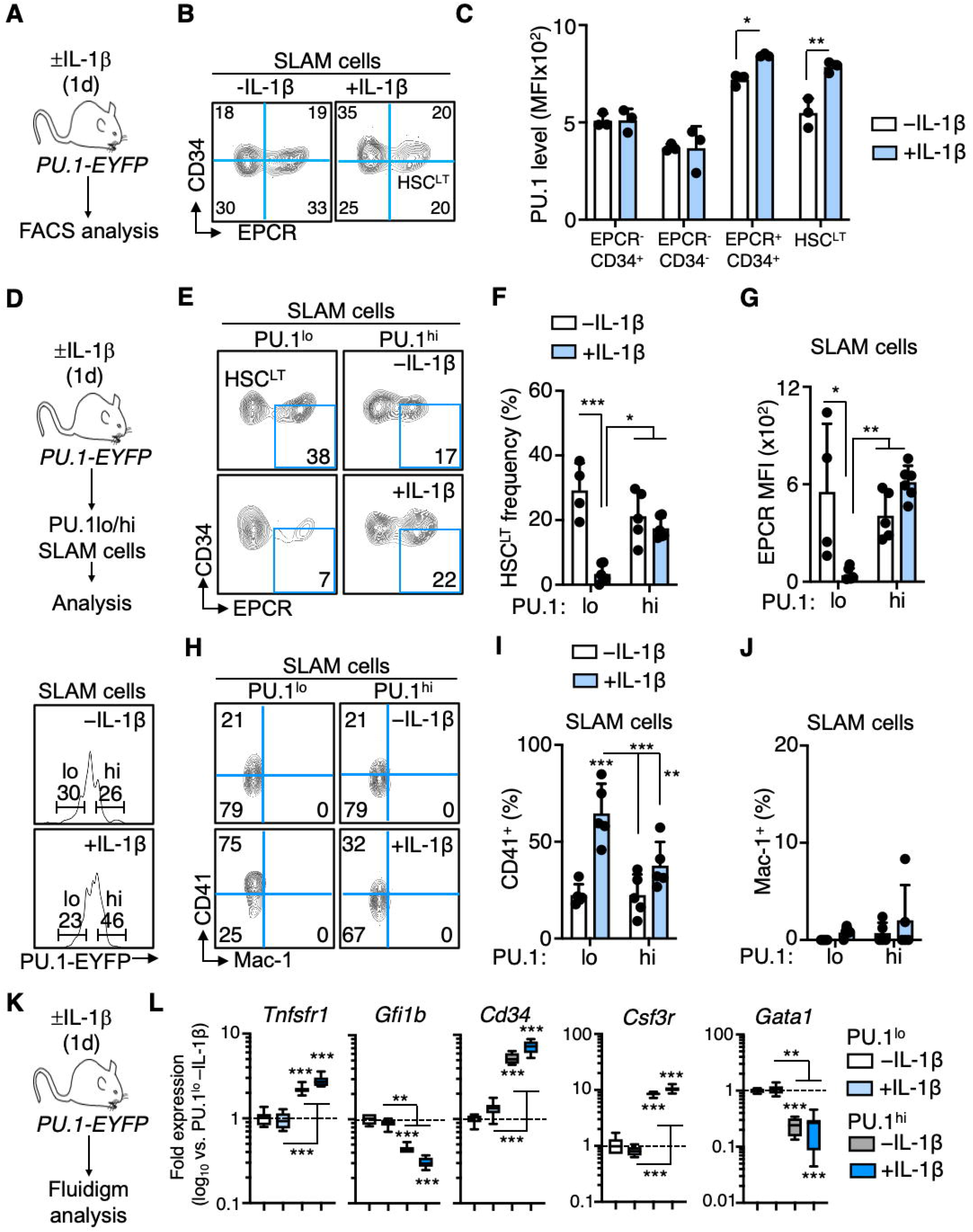
IL-1 rapidly triggers phenotypic remodeling of the PU.1^lo^ SLAM compartment. **A)** Experimental design for analysis of *PU*.*1-EYFP* mice treated for 1d ± IL-1 (n = 3/grp). **B)** Representative FACS plot showing fractionation of SLAM gate by EPCR and CD34 levels. **C)** Analysis of *PU*.*1-EYFP* reporter expression in SLAM fractions. Data are expressed as geometric mean fluorescence intensity (MFI). Individual values are shown, bars represent mean values. Data are representative of two independent experiments. **D)** Experimental design and representative FACS plots showing frequency of PU.1^hi^ and PU.1^lo^ SLAM cells, for analysis of PU.1^hi^ and PU.1^lo^ SLAM cells isolated from *PU*.*1-EYFP* mice treated for 1d ± IL-1 (n = 4-6/grp). **E)** Representative FACS plot showing phenotypic HSC^LT^ frequencies within the SLAM gate in *PU*.*1-EYFP* mice treated for 20d ± IL-1β. **F)** Summary data showing phenotypic HSC^LT^ frequency within the SLAM gate of *PU*.*1-EYFP* mice treated for 20d ± IL-1β (n = 4-6/grp). Individual values are shown, bars represent mean values. Data are compiled from two independent experiments. **G)** EPCR expression in phenotypic HSC^LT^ within the SLAM gate of *PU*.*1-EYFP* mice treated for 1d ± IL-1β (n = 4-6/grp). Individual values are shown, bars represent mean values. Data are compiled from two independent experiments. **H)** Representative FACS plot showing frequencies of CD41^+^ and Mac-1^+^ cells within the SLAM gate in *PU*.*1-EYFP* mice treated for 1d ± IL-1β. **I)** Frequency of CD41+ cells within the SLAM gate of *PU*.*1-EYFP* mice treated for 1d ± IL-1β (n = 5/grp). Individual values are shown, bars represent mean values. Data are representative of two independent experiments. **J)** Frequency of Mac-1+ cells within the SLAM gate of *PU*.*1-EYFP* mice treated for 1d ± IL-1β (n = 5/grp). Individual values are shown, bars represent mean values. Data are representative of two independent experiments. **K)** Experimental design for Fluidigm qRT-PCR array analysis of PU.1^lo^ and PU.1^hi^ SLAM cells from *PU*.*1-EYFP* mice treated for 1d ± IL-1β (n = 8/grp). **L)** Fluidigm qRT-PCR analysis of genes in PU.1^lo^ and PU.1^hi^ SLAM cells from *PU*.*1-EYFP* mice treated for 1d ± IL-1β. Data are expressed as log_10_ fold change versus PU.1^lo^ -IL-1β. Box represents upper and lower quartiles with line representing median value. Whiskers represent minimum and maximum values. Data are representative of two independent experiments. Data are shown as means ± SD. Significance was determined by ANOVA with Tukey’s post-test. *p<0.05; **p<0.01; ***p<0.001.

### Low PU.1 levels enrich for Mk activity in the SLAM compartment following IL-1 exposure

Our data suggest that following IL-1 exposure, the PU.1^lo^ SLAM compartment is rapidly enriched for cells that phenotypically resemble HSC-like Mk progenitors. To experimentally evaluate their functional potential, we isolated PU.1^lo^ and PU.1^hi^ SLAM cells from mice 1 day following injection with IL-1 or PBS and cultured them to assess lineage output (**Fig. 4A**). Consistent with our phenotypic data, microscopic examination of the cultures at day 3 revealed a rapid burst of Mk production from PU.1^lo^ SLAM cells, particularly those isolated from IL-1-treated mice (**Fig. 4B**). In support of our observation, PU.1^lo^ SLAM cultures were significantly enriched for CD41^+^ cells after 4 days of culture, again particularly in cultures of PU.1^lo^ SLAM cells isolated from IL-1-treated mice (**Fig. 4C-D**). On the other hand, PU.1^hi^ SLAM cell cultures contained a smaller fraction of CD41^+^ cells that was unaltered by *in vivo* IL-1 exposure. In line with their reduced repopulating activity following transplant and reduced phenotypic HSC^LT^ content, PU.1^lo^ SLAM cells isolated from IL-1-exposed mice also exhibited significantly reduced expansion in liquid culture (**Fig. S3A**). To confirm these results using a complementary approach and to measure impacts on short-term potential, we analyzed the clonogenic activity of PU.1^lo^ and PU.1^hi^ SLAM cells isolated from mice one day after injection with IL-1 or PBS (**Fig. 4E**). Consistent with our transplant data, PU.1^lo^ SLAM cells isolated from IL-1-injected mice had significantly reduced clonogenic activity (**Fig. 4F**). Likewise, in addition to the early burst of Mk activity in liquid culture, PU.1^lo^ SLAM cells from IL-1-injected mice retained an increased level of Mk potential, based on the proportion of Mk and Mix (myeloid + Mk) colonies present (**Fig. 4F**). On the other hand, the clonogenic activity and lineage distribution of PU.1^hi^ SLAM cells was essentially unchanged by *in vivo* IL-1 exposure and contained minimal Mk potential. Taken together, these data provide functional evidence that the PU.1^lo^ SLAM fraction consists largely of HSC-like progenitors with extensive Mk potential following IL-1 exposure.

**Figure 4:**
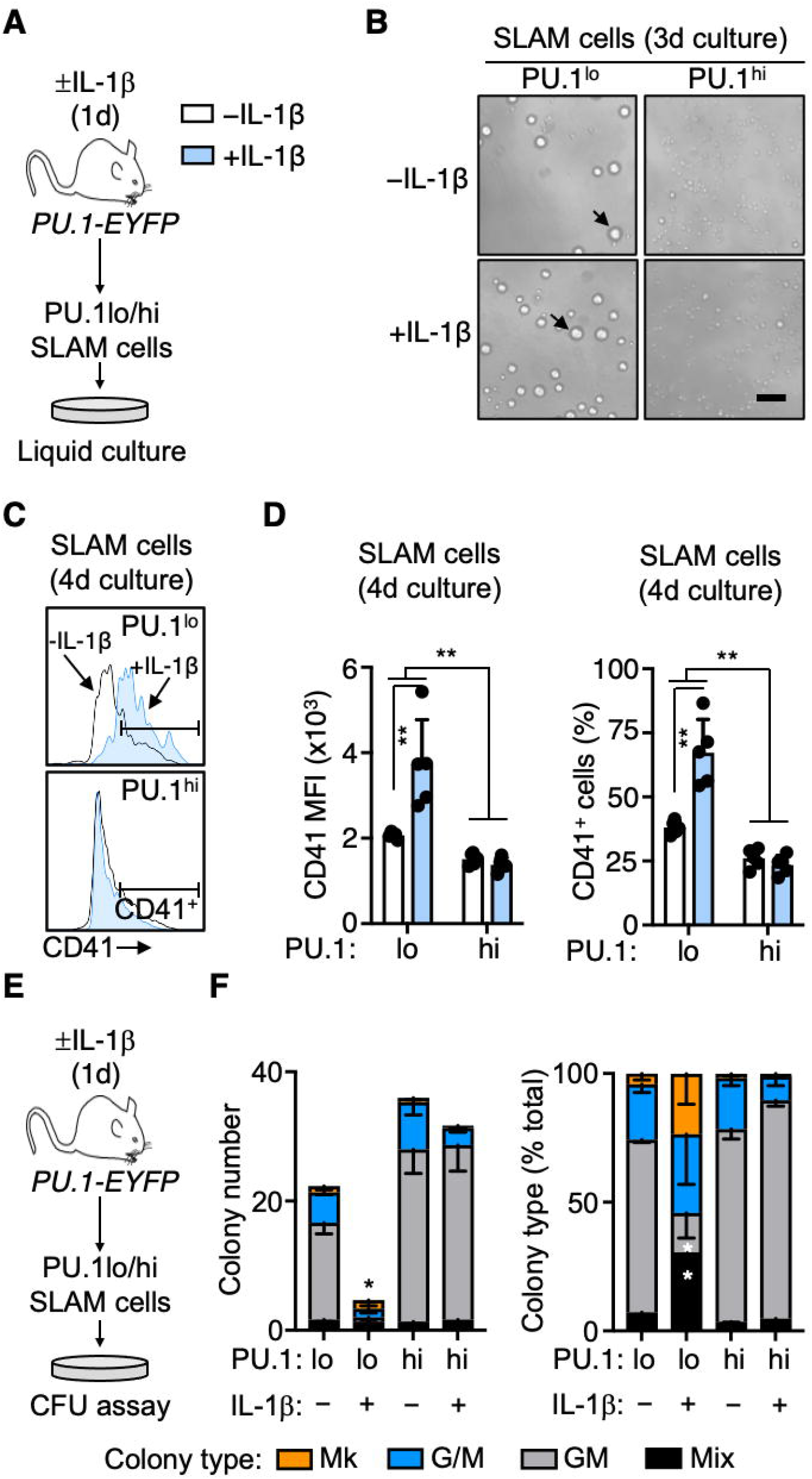
PU.1 levels identify Mk activity in the SLAM compartment following IL-1 exposure. **A)** Experimental design for *in vitro* liquid culture analysis of PU.1^lo^ and PU.1^hi^ SLAM cells from *PU*.*1-EYFP* mice treated for 1d ± IL-1β (n = 5/grp). **B)** Representative phase-contrast microscope images after 3 days culture of PU.1^lo^ and PU.1^hi^ SLAM cells from *PU*.*1-EYFP* mice treated for 1d ± IL-1β. Scale bar: 100μm. **C)** Representative histograms of CD41 expression after 4 days culture of PU.1^lo^ and PU.1^hi^ SLAM cells from *PU*.*1-EYFP* mice treated for 1d ± IL-1β. **D)** Summary data showing CD41 expression (left) and frequency of CD41^+^ cells (right) after 4 days culture of PU.1^lo^ and PU.1^hi^ SLAM cells from *PU*.*1-EYFP* mice treated for 1d ± IL-1β (n = 5/grp). Expression data are geometric mean fluorescence intensity (MFI). Individual values are shown, bars represent mean values. Data are representative of two independent experiments. **E)** Experimental design for *in vitro* methylcellulose culture analysis of PU.1^lo^ and PU.1^hi^ SLAM cells from *PU*.*1-EYFP* mice treated for 1d ± IL-1β (n = 3/grp). **F)** Colony number (left) and colony type distribution (right) of methylcellulose-cultured PU.1^lo^ and PU.1^hi^ SLAM cells from *PU*.*1-EYFP* mice treated for 1d ± IL-1β (n = 3/grp). Colony types were based on visual scoring. Colony type distribution is expressed as % total colonies. Data are representative of two independent experiments. Mk: megakaryocytic; G/M Data are shown as means ± SD. Significance was determined by ANOVA with Tukey’s post-test. *p<0.05; **p<0.01.

## Discussion

In the present study, we used functional assays, phenotypic analyses, and molecular profiling to resolve the relationship between PU.1 levels and HSC activity within the SLAM gate. We also set out to examine the extent to which inflammatory stress modulates these features. We find that PU.1 levels in the SLAM compartment define functionally and phenotypically heterogenous activities, with long-term HSC activity and phentotypic HSC^LT^ enriched in PU.1^lo^ SLAM cells at steady state but not in response to IL-1-induced inflammatory stress. Instead, IL-1 rapidly remodels the PU.1^lo^ SLAM fraction, which becomes enriched almost exclusively with EPCR^-^ HSC-like progenitors that possess extensive Mk potential but minimal repopulating and myeloid clonogenic activity. Altogether, our data demonstrate that PU.1 expression identifies functionally heterogenous and dynamically regulated activities within the phenotypic SLAM HSC compartment.

We previously found that the molecular profile of the PU.1^lo^ SLAM compartment changed significantly after IL-1 exposure, with reductions in HSC genes and concomitant increases in cell cycle activity^6^. Here, we find that the functional properties and cellular composition of the PU.1^lo^ SLAM compartment shift dramatically in the presence of IL-1. While HSC activity is clearly enriched in the PU.1^lo^ fraction at steady state, our transplantation and colony assays demonstrate that this activity is almost entirely lost following IL-1 exposure. Concurrently, we observe depletion of phenotypic EPCR^+^/CD34^-^ HSC^LT^ in PU.1^lo^ SLAM cells. These findings are consistent with our published observation that PU.1 is robustly induced in phenotypic HSC^LT^, which removes these cells from the PU.1^lo^ SLAM gate^6^. Thus, changes in gene expression and cell cycle status we observed previously in PU.1^lo^ SLAM cells are unlikely related to induction of a molecular program favoring proliferation and differentiation in HSC^LT^ themselves. Instead, they are caused by a shift in the composition of the PU.1^lo^ SLAM compartment. In line with these findings, we previously showed that the cell cycle activity and HSC gene expression profiles of different SLAM subsets identified based on coordinate expression of EPCR and CD34 are largely unchanged following chronic IL-1 exposure^10^. Thus, while low PU.1 levels are typically associated with long-term HSC activity, here we find that a heterogenous mix of activities in the SLAM gate are associated with low PU.1 expression. On the other hand, we find that the PU.1^hi^ gate exhibits an intermediate repopulating activity relative to steady-state PU.1^lo^ cells, regardless of IL-1 treatment. While the frequency of phenotypic HSC^LT^ (and even EPCR expression levels) is highly similar to HSC^LT^ in the PU.1^lo^ SLAM fraction at steady state, these cells appear to possess a more limited reconstitution capacity, perhaps owing to increased expression of myeloid lineage determinants that promote their differentiation. Taken together, these data reveal a striking degree of functional heterogeneity based on PU.1 expression within the SLAM compartment, particularly under inflammatory conditions.

Following IL-1 treatment, we find that the remaining PU.1^lo^ population is almost exclusively composed of EPCR^-^ SLAM cells that express high levels of CD41 and Mk/E lineage genes including *Gata1* and *Gfi1b*. These cells are reminiscent of HSC-like Mk progenitors that were previously characterized in the context of *in vivo* poly I:C treatment^17^. In that setting, CD41^+^ HSC-like Mk progenitors appeared to function as an ‘emergency’ stress-induced compartment that lacked multilineage reconstitution activity and rapidly gave rise to Mk lineage cells both *in vivo* and in culture. Similarly, we previously showed that the EPCR^-^ SLAM fraction lacks long-term repopulating activity and exhibits extensive Mk lineage priming, suggesting EPCR^-^ SLAM cells and HSC-like Mk progenitors are likely an overlapping cell population^10^. Consistent with these observations, PU.1^lo^ SLAM cells from IL-1-treated mice rapidly produce megakaryocytes and even retain high levels of CD41 expression in culture, in contrast to PU.1^hi^ cells that exhibit limited CD41 expression and Mk lineage potential in culture. Hence, the PU.1^lo^ SLAM fraction likely overlaps extensively with Vwf-GFP^+^ SLAM cells, which also enriches for HSC^LT^ at steady state but marks a predominantly EPCR^-^ SLAM cell population with limited reconstitution activity under inflammatory conditions^10^. It is worth noting that the ontological relationship between HSC^LT^ and EPCR^-^ SLAM cells is not well characterized. Specifically, it is not well understood whether EPCR^-^ SLAM cells represent a transient cellular state or the entry point into a terminal Mk differentiation pathway, either via direct differentiation or via MPP^MkE^ (MPP2)^20^. Previous analyses have shown that HSC-like Mk progenitors already exhibit Mk-like protein synthesis programs and an abundance of alpha granules in their cytoplasm, suggesting they have already adopted an immature Mk progenitor phenotype^17^. Furthermore, H2B-GFP labeling studies have shown that HSC with minimal divisional history retain an Mk gene program that is lost as they undergo successive cell divisions, suggesting that at least a subset of HSC^LT^ may rapidly adopt an HSC-like Mk progenitor state^21^. We also find that EPCR^-^ PU.1^lo^ SLAM cells possess a surface phenotype that is distinct from MkP, supporting a model^17^ in which these cells adopt a unique developmental trajectory allowing rapid platelet production in response to inflammatory stress. Thus, markers such as EPCR and CD41 can be used to readily separate HSC^LT^ activity from Mk progenitor activity in the phenotypic SLAM compartment, particularly under inflammatory stress conditions.

On the other hand, the PU.1^hi^ SLAM fraction exhibits limited Mk differentiation potential and CD41 expression. Instead, these cells express high levels of myeloid determinants such as Mac-1, *Gcsfr, Cd34* and *Cebpe* in response to inflammation, suggesting they may be primed for myeloid differentiation. We previously showed that IL-1-induced Mac-1 expression occurs exclusively in EPCR^+^ SLAM cells, in line with patterns of PU.1 expression shown here^6, 10^. It is worth noting however, that PU.1^hi^ cells do still exhibit limited expansion of CD41^+^ cells and at least some Mk activity *in vitro*, suggesting Mk activity is not completely extinguished by these increases in PU.1 expression. While classical models of hematopoietic lineage fate describe a cross-antagonistic relationship between PU.1 and Gata1 that leads to mutually exclusive myeloid/lymphoid versus Mk/E lineage fate decisions^22^, this model has been challenged by recent quantitative imaging studies showing that Gata1 and PU.1 can be stochastically expressed in the same cells^23^, and that Gata1 expression and subsequent Mk lineage commitment can occur in PU.1-expressing cells^24^. While our study was not designed to specifically resolve the relationship between PU.1 and Gata1 in specifying Mk versus granulocyte/macrophage lineage fates, our data do suggest that Mk priming is enriched in, but not necessarily exclusive to, PU.1^lo^ cells.

Altogether, our data show that the phenotypic SLAM compartment contains functionally heterogenous cell types defined by similar levels of PU.1 expression. While HSC^LT^ are nominally considered PU.1^lo^ cells, inflammatory signals induce PU.1 expression in these cells, separating out a parallel population of PU.1^lo^ HSC-like Mk progenitors with distinct molecular and functional attributes. The extent to which these HSC-like Mk progenitors are directly produced by HSC (and in particular are replenished by HSC^LT^ that adopt a PU.1^hi^ identity) remains to be elucidated, and future studies using clonal tracking approaches can be used to better define the hierarchical relationship between these different populations, as well as the rate at which HSC-like Mk progenitors are replenished by the HSC compartment, particularly under inflammatory stress. Lastly, our data argue for the adoption of rigorous approaches for prospectively isolating HSC^LT^ activity under non-steady state conditions and further assessment of transcriptional, metabolic and functional heterogeneity within HSC-enriched hematopoietic cell fractions.

## Materials and Methods

### Mice

Wild-type C57BL/6 and congenic B6.SJL-*Ptprc*^*a*^*Pepc*^*b*^/BoyJ (Boy/J) mice were obtained from The Jackson Laboratory. *PU*.*1-EYFP* mice^16, 25^ were a kind gift of Dr. Claus Nerlov (MRC Weatherall Institute). All animal procedures were approved by the University of Colorado Anschutz Medical Campus Institutional Animal Care and Use Committee (IACUC), protocol #00091. Mice were housed in a temperature- and light-controlled facility in HEPA-filtered cages and provided with chow and water *ad libitum*.

### Flow cytometry

Bone marrow harvest and flow cytometry were performed as previously described^6^. For sorting experiments, BM cells were isolated from the arms, legs, pelvis and spines by crushing bones in a mortar with a pestle containing staining media (SM--HBSS supplemented with 2% fetal bovine serum (FBS)) and transferred over a 70-micron filter. Cells were resuspended in ACK for five minutes, incubated on ice, and washed with SM. Erythrocyte depleted BM cells were resuspended in 3mL SM and transferred over a 70-micron filter. A ficoll gradient was created to separate out cellular debris by adding 2mL Histopaque 1119 (Sigma) to the bottom of the cell suspension followed by centrifugation at 1200 RPM (Beckman Coulter) with the break turned off. The top liquid phase containing BM cells was transferred into a new tube and washed once with SM. BM cells were resuspended in a solution of 100µL SM and 5µL murine CD117 microbreads (Miltenyi) per mouse, incubated on ice for 20 minutes, washed and enriched on an autoMACS Pro separator (Miltenyi). For immunophenotyping analyses, femurs and tibiae were flushed using a syringe containing 3mL SM and affixed with a 1 1/2 inch X 21-gauge needle. BM cells were erythrocyte depleted by ACK treatment, resuspended in SM, and counted using a ViCell automated cell-counter (Beckman Coulter). For HSPC analyses, BM cells were blocked with purified Rat IgG (Sigma) and stained with an antibody cocktail in SM containing CD3 PE/Cy5 (eBioscience, 15-0031-81), CD4 PE/Cy5 (eBioscience, 15-0041-82), CD5 PE/Cy5 (Biolegend, 100610), CD8 PE/Cy5 (Biolegend, 100710), B220 PE/Cy5 (Biolegend, 103210), Ter119 PE/Cy5 (Biolegend, 116210), Gr1 PE/Cy5 (Biolegend, 108410), EPCR PE (eBioscience, 12-2012-82), CD11b PE/Cy7 (Biolegend, 101216), CD34 AF647 (BD Biosciences, 560230), CD48 A700 (Biolegend, 103426), c-Kit APC/Cy7 (Biolegend, 105826), Flk2 Biotin (Biolegend, 135308), and incubated for 30 minutes on ice. Subsequently, cells were stained with an antibody cocktail in SM containing Brilliant Buffer (BD Biosciences), Sca1 Bv421 (Biolegend, 108128), CD41 BV510 (Biolegend, 113923), Streptavidin-BV605 (BD Biosciences, 563260), CD16/32 BV711 (Biolegend, 101337), and CD150 BV785 (Biolegend, 115937) for 30 minutes on ice. Cells were resuspended in a solution of SM containing Propidium Iodide (PI) for dead cell exclusion. For mature BM cell analyses to monitor BM chimerism, peripheral blood cells were RBC depleted using ACK as above and stained with an antibody cocktail of CD8-PE (Biolegend, 100708), Mac-1 PE/Cy7, IgM APC (eBioscience, 17-1590-82), CD3 AF700 (Biolegend, 100216), CD19 APC/Cy7 (Biolegend, 115530), Gr1 PB (Biolegend, 108430), CD4 BV510 (Biolegend, 100449), Ly6C BV605 (BD Biosciences, 563011), B220 BV875 (Biolegend, 103246), and incubated for 30 minutes on ice. Cells were washed and resuspended in PI. For liquid culture flow analyses, cells were stained with an antibody cocktail containing CD9 PE (Biolegend, 124805), Sca1 PE/Cy7 (Biolegend, 108114), CD11b APC (Biolegend, 101212), c-Kit APC/Cy7, Gr1 PB, CD41 BV510, CD16/32 BV711, CD150 BV785 and incubated on ice for 30 minutes. The cells were washed and resuspended in DPBS with PI. Data were acquired on a 12-channel, 3-laser FACSCelesta or an 18-channel, 5-laser LSRFortessa (Becton-Dickenson) and analyzed using FlowJo v10.

### *In vivo* procedures

*In vivo* IL-1β stimulation was performed as previously described^6^. Recombinant murine IL-1β was resuspended in sterile D-PBS/0.2% BSA and injected in a 100 μl bolus of 0.5 μg once daily via the intraperitoneal (i.p.) route, using a 31g insulin syringe. Transplantation experiments were performed as previously described^10^. CD45.1^+^ Boy/J recipient mice were lethally irradiated (11 Gy, split dose 2h apart) using a Cesium source (JL Shepherd & Associates, San Fernando, CA) and maintained on Bactrim in autoclaved water for 4 weeks post-transplant. Mice were injected retro-orbitally with 250 sorted SLAM cells (either PU.1^hi^ or PU.1^lo^) isolated from donor mice along with 5×10^5^ Boy/J helper cells that were Sca-1 depleted using an AutoMACS and resuspended in SM. Mice were bled retro-orbitally every four weeks for 20 weeks to monitor donor chimerism. Blood collection was performed via retro-orbital route into EDTA-coated microtainers. Complete Blood Count (CBC) analyses were performed on a calibrated HemaVet CBC analyzer.

### Cell culture

All cell cultures were performed in a humidified incubator (ThermoFisher Scientific) at 37°C and 5% CO_2_. For liquid and methylcellulose cultures, PU.1 eYFP low and high SLAM cells were purified from mice that were treated with either PBS or .5µg IL-1β (Peprotech) 24 hours prior to harvest. PU.1^lo/hi^ were selected based on median fluorescent intensity (MFI) where approximately the lowest 25% were designated PU.1^lo^ and the highest 25% designated as PU.1^hi^. Pools of 400 cells per well were sorted for liquid culture into Stempro 34 medium (Gibco, 10639011) and supplemented with antibiotic-antimycotic (Gibco, 15240062), 2mM L-glutamine (Gibco, 25030024), 25ng/mL IL-11 (Peprotech), 25ng/mL SCF (Peprotech), 25ng/mL TPO (Peprotech), 25ng/mL Flt3L (Peprotech), 10ng/mL IL-3 (Peprotech), 10ng/mL GM-CSF (Peprotech), and 4U/mL EPO (Peprotech). Half of the media was replenished every two days and cells were harvested for flow cytometry analysis and cell counts on days 4 and 8. Methylcellulose cultures were executed using Methocult M3231 Base Media (Stemcell Technologies, 03231) containing antibiotic-antimycotic, IMDM, 25ng/mL IL-11, 25ng/mL SCF, 25ng/mL TPO, 25ng/mL Flt3L, 10ng/mL IL-3, 10ng/mL GM-CSF, and 4U/mL EPO. Colonies were counted and phenotyped at day 8.

### Fluidigm qRT-PCR analysis

Fluidigm qRT-PCR analysis was performed as previously described^6^. Briefly, 100 PU.1 low and PU.1 high SLAM cells per well were sorted directly into 5µL of 2X Reaction Mix (Invitrogen) contained within a 96-well PCR plate. Once all cells were sorted, the plates were sealed with an aluminum seal, spun in a centrifuge at 1200 RPM for 5 minutes, snap frozen in liquid nitrogen, and stored at -80°C. cDNA was generated from RNA using Superscript III (Invitrogen) and a custom primer set mix (Fluidigm) that were amplified for 18 cycles on a thermocycler (Eppendorf). Excess primers were removed from the samples by Exonuclease I (NEB) treatment, and the samples were diluted in DNA suspension buffer (Teknova). Pre-amplified cDNA and custom primer sets were loaded onto a Fluidigm 96.96 Dynamic Gene Expression IFC. Subsequently, the IFC was run on a Biomark HD (Fluidigm) with SsoFast Sybr Green (Bio-Rad) used for detection. Fluidigm gene expression software was used to analyze the data and all values are relative to *Gusb*. Relative changes in gene expression were determined using the ΔΔCT approach.

### Statistical analysis

Statistical analyses were performed using GraphPad Prism. Multivariate comparisons were made using ANOVA with Tukey’s post-test. P-values <0.05 were considered statistically significant. For *in vivo* studies, groups of mice receiving treatments (IL-1, transplant) were chosen randomly within each cage. Experiments were repeated at least twice to ensure reproducibility of findings.

## Acknowledgements

We acknowledge Garrett Hedlund for expert flow cytometry assistance. This work was supported by R01 DK119394, the University of Colorado Outstanding Early Career Scholar Program and the Cleo Meador and George Ryland Scott Endowed Chair in Hematology (to E.M.P.), F31 HL138754 (to J.L.R.), the National Science Foundation Graduate Research Fellowship Program (NSF GFRP; to T.S.M.). K.E.N. was supported by a supplement to R01 DK119394. This work was supported in part by the University of Colorado Cancer Center Flow Cytometry Shared Resource, funded by NCI grant P30 CA046934.

## Author Contributions

Conceptualization: E.M.P. Methodology: E.M.P., J.L.R., J.S.C. Investigation: J.S.C., J.L.R., G.H.; T.S.M., K.E.N., P.D.C; Resources: E.M.P., P.D.C.; Writing - Original Draft: E.M.P., J.S.C.; Writing – Review/Editing: J.L.R., G.H., T.S.M., K.E.N., P.D.C.; Supervision: E.M.P.; Funding Acquisition: E.M.P.

## Supplementary Figure Legends

**Figure S1: CBC analysis and lineage distribution of transplant experiments**.

**A)** Complete blood count (CBC) analysis of blood from mice treated for 20d ± IL-1 (n = 9/grp). Means and individuals values are shown. Data are representative of two independent experiments.

**B)** Summary data showing donor myeloid (top) and lymphoid (bottom) lineage distribution in peripheral blood of transplanted recipient mice (n = 6-8/grp). Data are compiled from two independent experiments. Bars represent mean values. Data are shown as means ± SD. Significance was determined by ANOVA with Tukey’s post-test. *p<0.05; **p<0.01.

**Figure S2: Multiparameter FACS comparison of EPCR**^**-**^ **PU**.**1**^**lo**^ **SLAM cells and MkP**.

**A)** Experimental design (left) and representative gatings (right) for analysis of SLAM and MkP populations from GFP-Myc::PU.1-EYFP reporter mice treated for 20d ± IL-1β.

**B)** Representative FACS histograms with geometric mean fluorescence intensity (MFI; inset number) for each marker. Data are representative of at least three individual animals per condition.

**Figure S3: Growth kinetics of PU**.**1**^**lo**^ **and PU**.**1**^**hi**^ **SLAM cells in liquid culture**.

A) Summary data showing number of cells (expressed as log_10_) after 12 days culture of PU.1^lo^ and PU.1^hi^ SLAM cells from *PU*.*1-EYFP* mice treated for 1d ± IL-1β (n = 5/grp). Mean values are shown. Data are representative of two independent experiments.

Data are shown as means ± SD. Significance was determined by ANOVA with Tukey’s post-test.

*p<0.05; **p<0.01; ***p<0.001.

## References

1. Koschmieder, S.; Rosenbauer, F.; Steidl, U.; Owens, B. M.; Tenen, D. G., Role of transcription factors C/EBPalpha and PU.1 in normal hematopoiesis and leukemia. Int J Hematol 2005, 81 (5), 368–77.

2. Pang, S. H. M.; de Graaf, C. A.; Hilton, D. J.; Huntington, N. D.; Carotta, S.; Wu, L.; Nutt, S. L., PU.1 Is Required for the Developmental Progression of Multipotent Progenitors to Common Lymphoid Progenitors. Front Immunol 2018, 9, 1264.

3. Nutt, S. L.; Metcalf, D.; D’Amico, A.; Polli, M.; Wu, L., Dynamic regulation of PU.1 expression in multipotent hematopoietic progenitors. J Exp Med 2005, 201 (2), 221–31.

4. Dakic, A.; Wu, L.; Nutt, S. L., Is PU.1 a dosage-sensitive regulator of haemopoietic lineage commitment and leukaemogenesis? Trends Immunol 2007, 28 (3), 108–14.

5. Kueh, H. Y.; Champhekar, A.; Nutt, S. L.; Elowitz, M. B.; Rothenberg, E. V., Positive feedback between PU.1 and the cell cycle controls myeloid differentiation. Science 2013, 341 (6146), 670–3.

6. Chavez, J. S.; Rabe, J. L.; Loeffler, D.; Higa, K. C.; Hernandez, G.; Mills, T. S.; Ahmed, N.; Gessner, R. L.; Ke, Z.; Idler, B. M.; Nino, K. E.; Kim, H.; Myers, J. R.; Stevens, B. M.; Davizon-Castillo, P.; Jordan, C. T.; Nakajima, H.; Ashton, J.; Welner, R. S.; Schroeder, T.; DeGregori, J.; Pietras, E. M., PU.1 enforces quiescence and limits hematopoietic stem cell expansion during inflammatory stress. J Exp Med 2021, 18 (6).

7. Solomon, L. A.; Podder, S.; He, J.; Jackson-Chornenki, N. L.; Gibson, K.; Ziliotto, R. G.; Rhee, J.; DeKoter, R. P., Coordination of Myeloid Differentiation with Reduced Cell Cycle Progression by PU.1 Induction of MicroRNAs Targeting Cell Cycle Regulators and Lipid Anabolism. Mol Cell Biol 2017, 37 (10).

8. Singh, H.; DeKoter, R. P.; Walsh, J. C., PU.1, a shared transcriptional regulator of lymphoid and myeloid cell fates. Cold Spring Harb Symp Quant Biol 1999, 64, 13–20.

9. Staber, P. B.; Zhang, P.; Ye, M.; Welner, R. S.; Nombela-Arrieta, C.; Bach, C.; Kerenyi, M.; Bartholdy, B. A.; Zhang, H.; Alberich-Jorda, M.; Lee, S.; Yang, H.; Ng, F.; Zhang, J.; Leddin, M.; Silberstein, L. E.; Hoefler, G.; Orkin, S. H.; Gottgens, B.; Rosenbauer, F.; Huang, G.; Tenen, D. G., Sustained PU.1 levels balance cell-cycle regulators to prevent exhaustion of adult hematopoietic stem cells. Mol Cell 2013, 49 (5), 934–46.

10. Rabe, J. L.; Hernandez, G.; Chavez, J. S.; Mills, T. S.; Nerlov, C.; Pietras, E. M., CD34 and EPCR coordinately enrich functional murine hematopoietic stem cells under normal and inflammatory conditions. Exp Hematol 2020, 81, 1–15 e6.

11. Pietras, E. M.; Mirantes-Barbeito, C.; Fong, S.; Loeffler, D.; Kovtonyuk, L. V.; Zhang, S.; Lakshminarasimhan, R.; Chin, C. P.; Techner, J. M.; Will, B.; Nerlov, C.; Steidl, U.; Manz, M. G.; Schroeder, T.; Passegue, E., Chronic interleukin-1 exposure drives haematopoietic stem cells towards precocious myeloid differentiation at the expense of self-renewal. Nat Cell Biol 2016, 18 (6), 607–18.

12. Pietras, E. M., Inflammation: a key regulator of hematopoietic stem cell fate in health and disease. Blood 2017, 130 (15), 1693–1698.

13. Etzrodt, M.; Ahmed, N.; Hoppe, P. S.; Loeffler, D.; Skylaki, S.; Hilsenbeck, O.; Kokkaliaris, K. D.; Kaltenbach, H. M.; Stelling, J.; Nerlov, C.; Schroeder, T., Inflammatory signals directly instruct PU.1 in HSCs via TNF. Blood 2019, 133 (8), 816–819.

14. Yamashita, M.; Passegue, E., TNF-alpha Coordinates Hematopoietic Stem Cell Survival and Myeloid Regeneration. Cell Stem Cell 2019, 25 (3), 357–372 e7.

15. Kiel, M. J.; Yilmaz, O. H.; Iwashita, T.; Yilmaz, O. H.; Terhorst, C.; Morrison, S. J., SLAM family receptors distinguish hematopoietic stem and progenitor cells and reveal endothelial niches for stem cells. Cell 2005, 121 (7), 1109–21.

16. Hoppe, P. S.; Schwarzfischer, M.; Loeffler, D.; Kokkaliaris, K. D.; Hilsenbeck, O.; Moritz, N.; Endele, M.; Filipczyk, A.; Gambardella, A.; Ahmed, N.; Etzrodt, M.; Coutu, D. L.; Rieger, M. A.; Marr, C.; Strasser, M. K.; Schauberger, B.; Burtscher, I.; Ermakova, O.; Burger, A.; Lickert, H.; Nerlov, C.; Theis, F. J.; Schroeder, T., Early myeloid lineage choice is not initiated by random PU.1 to GATA1 protein ratios. Nature 2016, 535 (7611), 299–302.

17. Haas, S.; Hansson, J.; Klimmeck, D.; Loeffler, D.; Velten, L.; Uckelmann, H.; Wurzer, S.; Prendergast, A. M.; Schnell, A.; Hexel, K.; Santarella-Mellwig, R.; Blaszkiewicz, S.; Kuck, A.; Geiger, H.; Milsom, M. D.; Steinmetz, L. M.; Schroeder, T.; Trumpp, A.; Krijgsveld, J.; Essers, M. A., Inflammation-Induced Emergency Megakaryopoiesis Driven by Hematopoietic Stem Cell-like Megakaryocyte Progenitors. Cell Stem Cell 2015, 17 (4), 422–34.

18. Pronk, C. J.; Rossi, D. J.; Mansson, R.; Attema, J. L.; Norddahl, G. L.; Chan, C. K.; Sigvardsson, M.; Weissman, I. L.; Bryder, D., Elucidation of the phenotypic, functional, and molecular topography of a myeloerythroid progenitor cell hierarchy. Cell Stem Cell 2007, 1 (4), 428–42.

19. van der Meer, L. T.; Jansen, J. H.; van der Reijden, B. A., Gfi1 and Gfi1b: key regulators of hematopoiesis. Leukemia 2010, 24 (11), 1834–43.

20. Challen, G. A.; Pietras, E. M.; Wallscheid, N. C.; Signer, R. A. J., The Simplified MPP Isolation Scheme: establishing a consensus approach for multipotent progenitor identification. Exp Hematol 2021.

21. Bernitz, J. M.; Kim, H. S.; MacArthur, B.; Sieburg, H.; Moore, K., Hematopoietic Stem Cells Count and Remember Self-Renewal Divisions. Cell 2016, 167 (5), 1296–1309 e10.

22. Burda, P.; Laslo, P.; Stopka, T., The role of PU.1 and GATA-1 transcription factors during normal and leukemogenic hematopoiesis. Leukemia 2010, 24 (7), 1249–57.

23. Wheat, J. C.; Sella, Y.; Willcockson, M.; Skoultchi, A. I.; Bergman, A.; Singer, R. H.; Steidl, U., Single-molecule imaging of transcription dynamics in somatic stem cells. Nature 2020, 583 (7816), 431–436.

24. Strasser, M. K.; Hoppe, P. S.; Loeffler, D.; Kokkaliaris, K. D.; Schroeder, T.; Theis, F. J.; Marr, C., Lineage marker synchrony in hematopoietic genealogies refutes the PU.1/GATA1 toggle switch paradigm. Nat Commun 2018, 9 (1), 2697.

25. Kirstetter, P.; Anderson, K.; Porse, B. T.; Jacobsen, S. E.; Nerlov, C., Activation of the canonical Wnt pathway leads to loss of hematopoietic stem cell repopulation and multilineage differentiation block. Nat Immunol 2006, 7 (10), 1048–56.

