## Supplementary figures and images for "PU.1 expression defines distinct functional activities in the phenotypic HSC compartment in a mouse model of inflammatory stress"

### Supplemental Figures

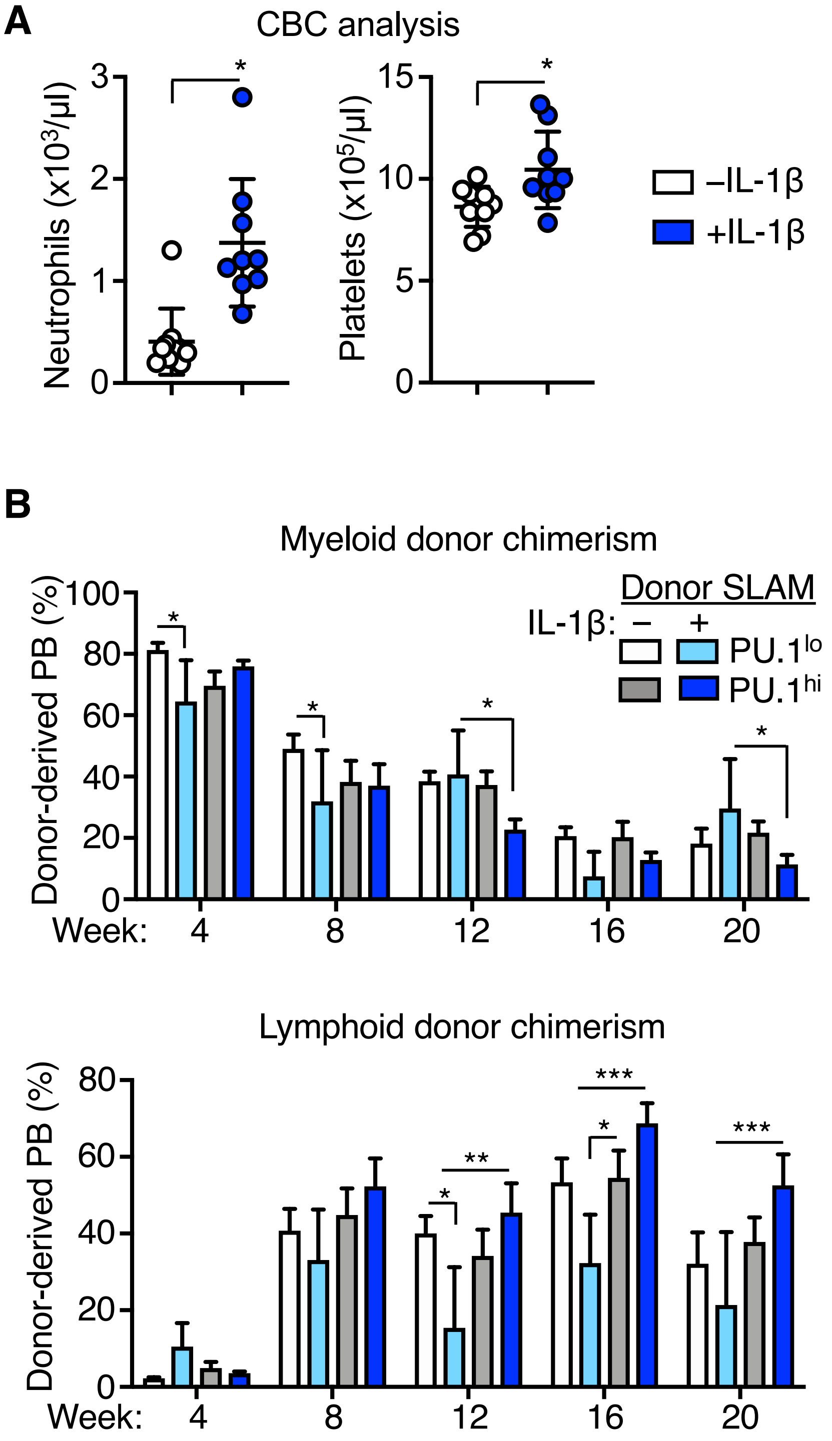

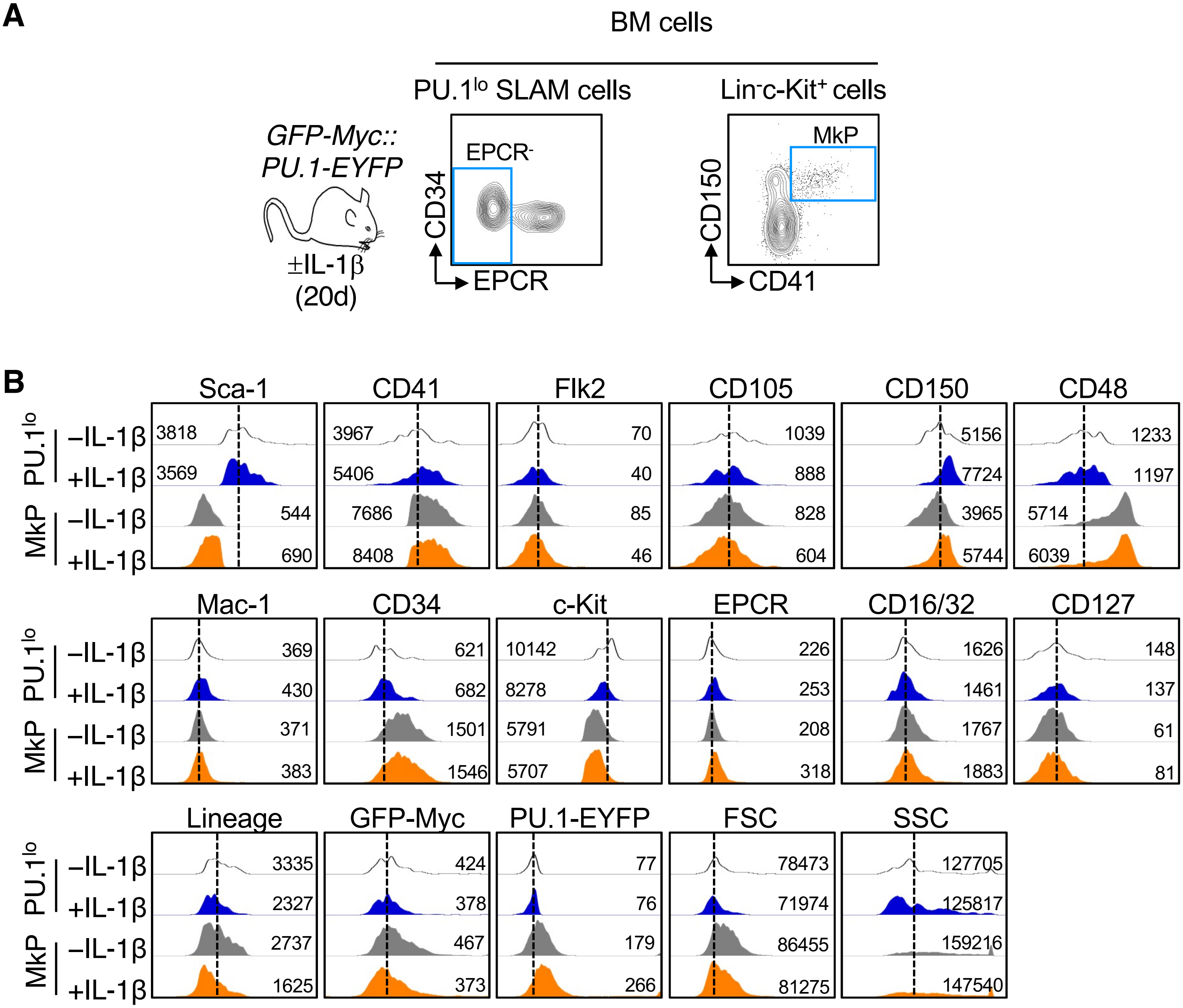

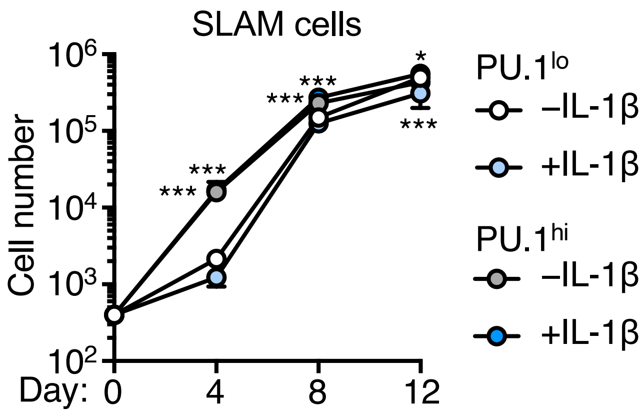
